# Monkeys Predict US Elections

**DOI:** 10.1101/2024.09.17.613526

**Authors:** Yaoguang Jiang, Annamarie Huttunen, Naz Belkaya, Michael L Platt

## Abstract

How people vote often defies rational explanation. Physical traits sometimes sway voters more than policies do–but why? Here we show that rhesus macaques, who have no knowledge about political candidates or their policies, implicitly predict the outcomes of U.S. gubernatorial and senatorial elections based solely on visual features. Given a pair of candidate photos, monkeys spent more time looking at the loser than the winner, and this gaze bias predicted not only binary election outcomes but also the candidates’ vote share. Analysis of facial features revealed candidates with more masculine faces were more likely to win an election, and vote share was a linear function of jaw prominence. Our findings endorse the idea that voters spontaneously respond to evolutionarily conserved visual cues to physical prowess and that voting behavior is shaped, in part, by ancestral adaptations shared with nonhuman primates.

**Significance Statement:** We report that monkeys and humans alike respond spontaneously to evolutionarily conserved facial masculinity cues in political candidates, and this innate sensitivity partly shapes voting behavior, highlighting the imperative for voters to overcome this ancient heuristic by becoming more informed on candidates and their policies.

## Introduction

Former US President Bill Clinton once remarked that “Americans prefer strong and wrong over weak and right”, highlighting the fact that voters sometimes choose to the detriment of their own economic wellbeing and personal liberty (1-3). Evidence-based deliberation by well-informed voters is a bedrock assumption of a functioning democracy (4, 5), so understanding, and potentially mitigating, deviations from rational decision-making by voters is critical for the health of our body politic.

Beyond politics, psychologists and behavioral economists have documented an extensive array of “irrational” decision-making tendencies, including loss aversion, sunk cost fallacy, the endowment effect, the “tyranny of choice,” and the disposition effect (6, 7). Such deviations from rational decision making have been hypothesized to reflect engagement of so-called “System 1” processing, which is fast, unconscious, intuitive, and emotional, by contrast with “System 2,” which is slow, conscious, deliberate, and cognitive (8). Recent work in behavioral economics and neuroeconomics (9-12) proffers that neural circuits driving decisions are optimized to obey constraints and solve problems confronted earlier in evolution–a perspective known as “ecological rationality” (12). The impacts of such deeply baked-in design principles in our neurobiology are not limited to voting behavior and microeconomics, but extend to consumer behavior, corporate culture, healthcare, domestic policy-making, international relations, and jurisprudence (7,8, 12–14).

Ecological rationality may help explain why the physical appearance of a candidate for office impacts their electability (15). Todorov and colleagues showed that people can accurately predict election outcomes based on very short visual exposure to candidate photos alone (16-18), including unfamiliar elections in foreign countries (19, 20). Remarkably, even preschool children can predict with above chance accuracy who will win an election based solely on candidate photos (21). In these studies, participants were asked to judge the competence, trustworthiness, and warmth of candidates, with competence judgments best predicting election results (16–18, 22). Participants further expressed the belief that competence is one of the most important attributes for politicians (16, 22, 23). These findings led Todorov and colleagues to propose a dual-process framework, in which voters assess candidate competence, partially on the basis of visual appearance, and cast votes accordingly.

While abundant evidence supports the idea that people quickly form opinions of others based on first impressions, which include visual appearance (24), the mechanism underlying these biases remains unclear, and the reasons why such biases persist in contemporary society remain obscure. Here we test the hypothesis that masculinized facial features, such as wide jaws and less prominent cheekbones, contribute to election outcomes, and that the impacts of physical cues to social dominance on decision making are evolutionarily conserved. To do so, we build upon previous research showing children can predict election outcomes based on candidate pictures alone (21), and extend it by testing whether macaque monkeys (*Macaca mulatta*) spontaneously do so as well. Prior research shows macaques attend to faces (25, 26), trade alimentary rewards for brief glimpses of dominant monkeys (27, 28), preferentially follow the gaze of dominant monkeys (29), and fixate longer on the faces of subordinate and female rather than dominant male monkeys (30, 31). Thus, nonhuman primates prioritize visual social information and are sensitive to cues associated with social dominance (32).

In the current study, monkeys freely examined pairs of candidate photos from prior U.S. senatorial, gubernatorial, and presidential elections. They displayed a consistent bias to look at losing candidates over winning candidates. “Votes” calculated from monkey gaze biases predicted election results as well as humans did based solely on visual information, and they were linearly correlated with vote share. To explore the contribution of masculinized facial features to this behavioral pattern, we measured the width of the jaw relative to the cheekbone in each candidate photograph and found their ratio predicted monkey gaze behavior and, moreover, linearly forecast vote share in US elections. Our findings endorse the idea that voters instinctively respond to evolutionarily conserved visual cues to physical prowess and masculinity, and that voting behavior is shaped, in part, by ancestral adaptations shared with nonhuman primates.

## Results

We examined the spontaneous gaze patterns of male macaques presented with pairs of candidate photos drawn from U.S. senatorial and gubernatorial elections (Figure 1a). To minimize familiarity effects on gaze behavior, in each session each pair of candidates was displayed only once. Each monkey (n = 3) participated in 2 sessions, with an inter-session interval of at least 3 days (n = 6 sessions total). All monkeys looked more at loser rather than winner pictures (Figure 1a), sometimes avoiding the winner altogether (Supplementary Figure 1a). Similarly, monkeys showed biased gaze towards female over male candidates (Figure 1a bottom, Supplementary Figure 1a bottom). Both biases were significant for all monkeys combined (Figure 1b, loser vs winner: P < 0.001, Wilcoxon signed rank; female vs male: P < 0.001, Wilcoxon rank sum), and consistent across the 3 subject monkeys (Supplementary Figure 1b).

**Figure 1.**
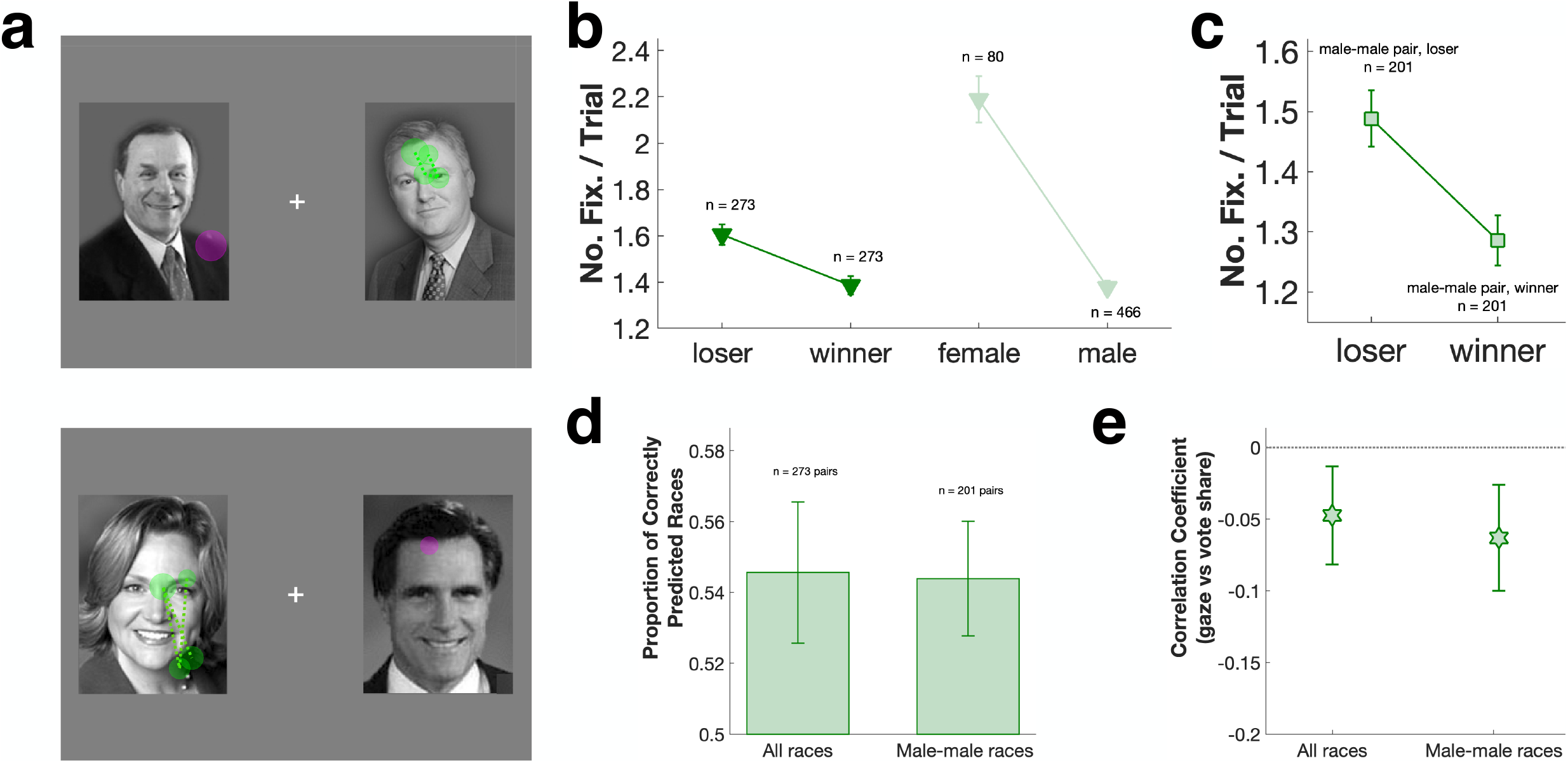
**a:** Example gaze patterns for gubernatorial elections in Oregon (2006, top) and Massachusetts (2002, bottom). Cross: central fixation spot. Filled circles: fixations on candidate pictures; circle sizes: fixation durations. Magenta: fixations on election winner; green: fixations on loser. **b:** Monkey gaze bias as a function of election outcome and candidate gender. **c:** Gaze bias restricted to male-male races. **d:** Average proportion of elections correctly predicted by gaze bias (n = 3 monkeys, 6 gubernatorial and 6 senatorial sessions in total). Error bars: mean ± SEM. **e:** Correlation of number of fixations on candidate with vote share. Error bars: 95% confidence interval.

Gaze biases towards losing candidates and female candidates interacted when a female ran against a male (n = 66 pairs making up 24.2% of all the races). Monkeys showed the strongest gaze bias towards female candidates who lost to a male (Figure 1a bottom, Supplementary Figure 1a bottom). When a female candidate beat a male opponent (Supplementary Figure 1c), however, the loser bias was sometimes nullified (Supplementary Figure 1c bottom, Supplementary Figure 1d right). By contrast, for male-male candidate pairs (n = 201 pairs making up 73.6% of all the races) the loser gaze bias was linear and strong (P = 0.002, Wilcoxon signed rank, Figure 1c).

We used monkeys’ gaze bias as ‘votes’ and compared them to real election outcomes. As Todorov and colleagues showed in humans (16-18), monkeys predicted U.S. gubernatorial and senatorial elections with above chance accuracy, for all races (54.6 ± 2.0%, P = 0.043, t-test, Figure 1d left) as well as male-male races only (54.4 ± 1.6%, P = 0.020, t-test, Figure 1d right). These patterns remained robust whether monkey ‘votes’ were tallied based on number of fixations (as in Figure 1d), fixation durations, or first fixation target (Supplementary Figure 1e). Monkey’s gaze bias also significantly predicted vote share for each candidate (number of fixations vs. vote share, all races: r = −0.048, P = 0.007; male-male races: r = −0.061, P = 0.003, Figure 1e). Human voters typically demonstrate a strong preference for incumbents (33, 34), but the correlation between monkey gaze bias and vote share was maintained against this incumbent effect (Supplementary Figure 1f).

We assume monkeys were unaware of the identity, party affiliation, or policies of any of the candidates. The only information accessible to them was candidate photos, suggesting monkeys predicted election outcomes and vote share based on visual features. Prior studies have shown monkeys are vigilant to information about social dominance: they will pay for a brief glimpse of a dominant male, but then rapidly look away to avoid direct eye contact, which is considered a sign of aggression (27, 28, 30, 31, 35-37). Due to their viewing bias towards female candidates, we surmised that monkeys were sensitive to masculine facial features, which are shared by monkeys and humans and associated with dominant social status–which for male macaques is correlated with physical prowess, peaks in adulthood, and declines in older age (37). For both monkeys and humans, compared with females, males on average have narrower cheekbones, wider jaws, higher facial width to height ratio (FWHR), and higher lower face prominence (LFP), resulting from high circulating testosterone during puberty (38-40). We measured all these features in candidate photos (Figure 2a), and found that jaw prominence–the ratio of jaw width to cheekbone width–accounted for 7-8% of the variance in vote share (gubernatorial races: r = 0.26, P < 0.001, senatorial races: r = 0.29, P < 0.001, overall: r = 0.27, P < 0.001, Figures 2b-c). On average, the winning candidate’s jaw was 2% more prominent than the losing candidate’s jaw (Supplementary Figure 2a). This pattern was consistent across gubernatorial and senatorial races, party affiliation, and incumbency (Supplementary Figure 2b). In alignment with previous literature, jaw prominence, cheekbone width, jaw width, FWHR, and LFP all reliably distinguished males from females in our sample of candidate photos (Supplementary Figure 2c). Only jaw prominence, however, distinguished faces of winners and losers (Supplementary Figure 2d).

**Figure 2.**
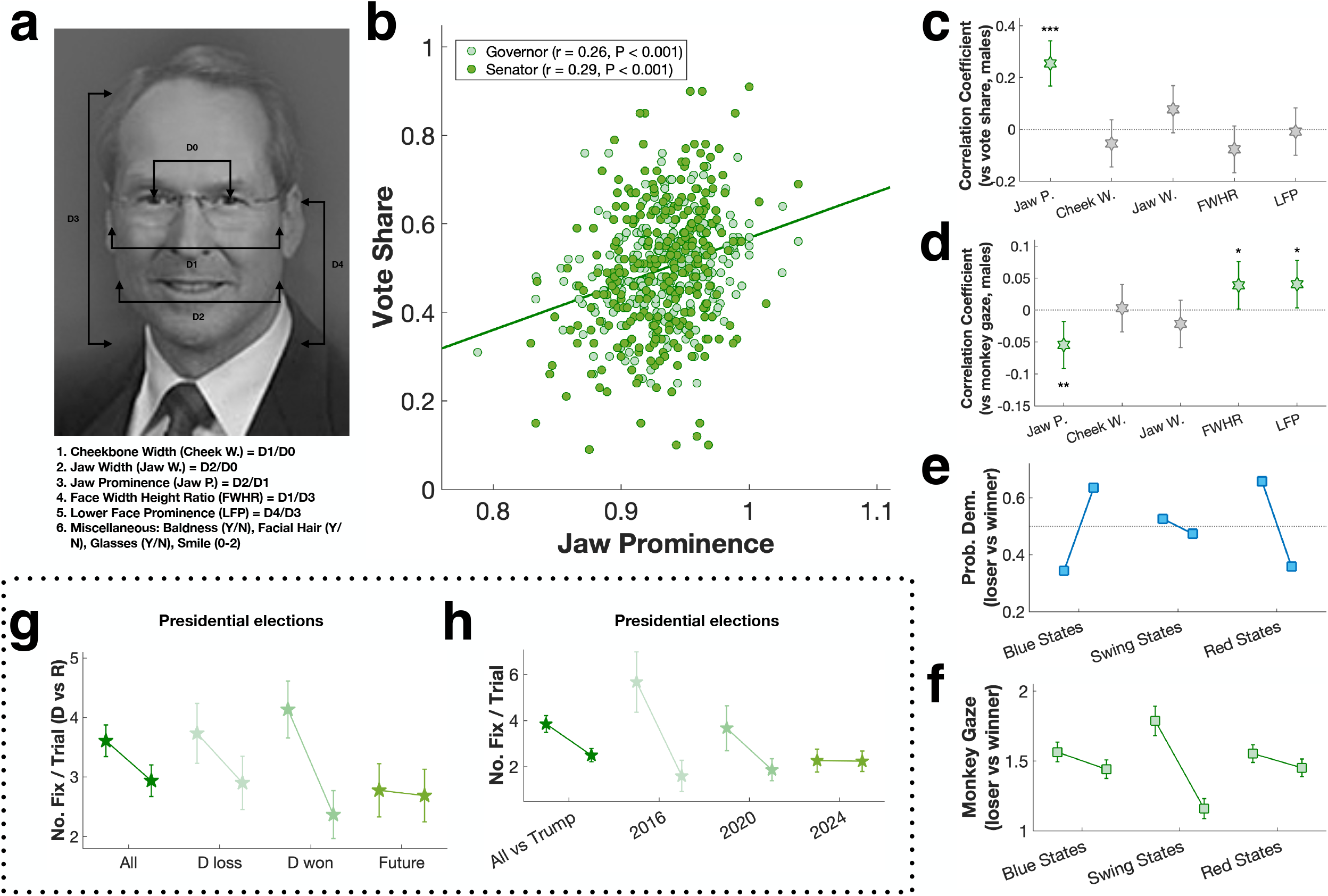
**a:** Facial feature measurements. **b:** Correlation of jaw prominence (jaw width/cheekbone width) with vote share. Line: linear regression. **c:** Correlations between all masculine facial features and vote share. **d:** Correlation of monkey gaze bias and vote share. Error bars: 95% confidence interval. **e:** The probability of the Democratic candidate winning versus losing in blue states, swing states, and red states. **f:** Monkey gaze bias for blue state, swing state, and red state elections. **g:** Gaze bias for Democratic and Republican presidential candidates as a function of election outcome. **h:** Gaze bias for elections between Donald Trump and his Democratic opponents over three successive elections. Error bars: mean ± SEM.

Jaw prominence also significantly predicted monkey’s gaze bias (r = −0.05, P = 0.004, Figure 2d)—the more prominent the jawline, the more likely monkeys were to avert their gaze. FWHR and LFP also weakly predicted monkey gaze, but in the opposite direction (r = 0.04, P = 0.041, and r = 0.04, P = 0.032, respectively, Figure 2d). In other words, candidates with the most prominent jaws, relative to their cheekbones, received the most votes, and monkeys most strongly avoided looking at them. Taken together, these findings suggest visual cues to physical prowess shape both visual orienting by monkeys and the choices humans make in the voting booth.

Abundant evidence shows competence judgments shape voting behavior (16-18). In prior studies, the relationship between competence judgements and voting persisted after controlling for perceived masculinity (22). We found jaw prominence was significantly correlated with competence rating (overall: r = 0.14, P = 0.001, males only: r = 0.12, P = 0.012, Supplementary Figures 2e). Partial correlation analyses revealed, and a general linear model confirmed, that facial masculinity and perceived competence each uniquely contributed to vote share (competence vs vote share: r = 0.31. P < 0.001; jaw prominence vs vote share: r = 0.25, P < 0.001). When combined, these two factors, based solely on photos of candidate faces, explained more than 15% of the variance in vote share.

Gaze bias in monkeys and voting by humans diverge, however, for female and older candidates. Overall, we found similar relationships between jaw prominence, competence rating, and vote share for female candidates as for males. That is, female candidates with more prominent jaws were rated as more competent (r = 0.30, P = 0.010, Supplementary Figure 2e, right), and also received more votes (r = 0.39, P < 0.001, Supplementary Figure 2f, left). Monkeys, by contrast, were biased to view female over male pictures, thereby weakening their predictions of races between females and males (Supplementary Figure 1d, Supplementary Figure 2g). Similarly, monkeys were biased to look at older rather than younger candidates (r = 0.06, P = 0.001), whereas human voters typically prefer older candidates (age vs vote share: r = 0.24, P < 0.001 in our sample, Supplementary Figure 2h; 22). We found no relationship between age and jaw prominence (r = 0.00, P = 0.976), or age and competence (r = 0.05, P = 0.261), indicating that age represents a dimension independent of masculinity and competence.

In addition to female and older candidates, human voters also chose challengers (as opposed to incumbents) and Democrats (as opposed to Republicans) more often than monkeys predicted they would (Supplementary Figure 3a). Party affiliation is a major predictor of voting behavior (41), so we examined how well monkeys’ gaze biases corresponded with voter behavior in blue (Democratic-leaning), red (Republican-leaning), and swing states (Figure 2e). We found that monkey gaze behavior predicted swing-state election results best (Figure 2f), and swing-state elections were most sensitive to facial masculinity cues (Supplementary Figures 3b-c).

Finally, we attempted to predict the upcoming 2024 US presidential election. For past presidential elections (2000-2020), monkeys’ predictions were at chance (50.4 ± 5.7%, P = 0.943, t-test), possibly reflecting voters’ recent preferences for Democratic and older candidates. As observed in the gubernatorial and senatorial elections, monkeys were biased to look at the Democratic over the Republican presidential candidate regardless of election outcome (P = 0.042, Wilcoxon signed rank; Figure 2g). This pattern was consistent across all 3 monkeys (Supplementary Figure 3d). As for the most recent races involving Donald Trump (2016-2024), monkeys’ gaze bias most strongly differentiated Trump from Hillary Clinton, less so from Joe Biden, and not at all from Kamala Harris (Supplementary Figure 3e). Thus, among the 3 most recent democratic nominees, based solely on visual features, Harris would be predicted to stand the best chance of winning, possibly reflecting Trump’s advanced age (Figure 2h) or voters detecting qualities in Trump other than physical dominance—for example low warmth, honesty, or likeability—that are deemed undesirable for politicians, as well as familiarity with his character and past performance, thus accounting for his vote share in 2020 underperforming predictions based on his jaw prominence (Supplementary Figure 3f).

## Discussion

Here we show for the first time that forecasts based on the gaze behavior of monkeys predict the results of U.S. gubernatorial and senatorial races–and do so as well as human adults and children do. Sensitivity to facial masculinity, particularly jaw prominence relative to cheekbones, best explained monkeys’ gaze bias. We surmise that people tested in prior studies similarly predicted election outcomes based on visual cues to physical prowess and dominance, and that vigilance for these features shapes voter choices in real elections.

Ecological rationality provides a potential explanation for the impact of facial masculinity on voting behavior (12, 13). Masculine facial features, such as wide jaw and prominent lower face are associated with high testosterone (38-40). Macaque monkeys can infer not only identity and reproductive state from conspecific faces, but also social status, indicating sensitivity to facial masculinity (25, 42, 43). Humans can also detect facial masculinity, and consider more masculine male faces to be not only more attractive (38-40, 44, 45) but also more likely to succeed (46). Mechanistically, human and nonhuman primates share brain regions and networks that prioritize visual social information with adaptive value (32, 47, 48). We hypothesize that in both humans and monkeys these shared structural and functional specializations spontaneously detect facial masculinity in candidates, thereby shaping attention and downstream processing, which, in humans, ultimately impacts voting behavior.

We note that apparent deep homologies in social attention shared by human and nonhuman primates do not explain all contemporary voting behavior. Based solely on facial masculinity cues, female candidates are projected to lose most races. Yet voters chose the female candidate about half the time (overall female winning probability = 48.8% in our sample), indicating other factors besides facial masculinity contribute to voting decisions. Similarly, voters preferred older candidates, although, based on gaze bias, monkeys found them less dominant looking. Voters also selected challengers and Democratic candidates more often than predicted by monkeys. These findings indicate factors beyond perceived masculinity and physical prowess shape voting behavior.

Our findings have implications for political campaigns. For example, judgment of facial masculinity relies on visual perception of low-spatial frequency information such as jaw width and cheekbone width, which can occur during brief presentations of low resolution images, such as peripheral glimpses of a campaign flier, thus supporting print media campaigns. Further, campaigns can strategically use photos emphasizing or de-emphasizing jaw width and cheekbone width (49, 50), which may explain why most candidates smile in their official portraits (92.3% of candidates smiled in the photos in our sample), thereby accentuating the jawline. Choices also depend on how a decision is framed (6-8). For example, Little et al. (51, Study 2) created morphed faces of George W. Bush and John Kerry, and participants were asked to indicate which one they would “vote for to run your country” in a time of peace or a time of war. In peacetime, the more Kerry-shaped face was preferred, but in wartime the more Bush-shaped face was preferred. The more Bush-shaped face was judged to be more masculine and dominant, traits participants favored in a time of conflict, but less intelligent and forgiving, traits participants favored in peacetime. Campaigns can frame social issues to their advantage by emphasizing external threats if the candidate is more masculine looking, or domestic tranquility if the candidate is more feminine looking, which may explain the apparent effectiveness of the Harris-Walz campaign’s focus on joy. Overall, our findings compel the development of strategies that encourage voters to become well-informed about candidates and their policies to help overcome evolutionarily ancient heuristics prioritizing visual predictors of physical prowess and masculinity.

## Material and Methods

### Animals

All procedures reported in this study were approved by the Institutional Animal Care and Use Committee of the University of Pennsylvania, and performed in accordance with *The Guide to the Care and Use of Laboratory Animals*. Three male rhesus macaques (M1: C, 15 years old, 15 kg; M2: L, 16 years old, 11 kg; M3: O, 17 years old, 17 kg) participated in the senatorial and gubernatorial election experiment, each for 2 days/sessions, with an intersession interval of at least 3 days (n = 6 sessions in total). Stimulus sets for each session consisted of 124 pairs of gubernatorial candidate pictures and 149 pairs of senatorial candidate pictures (n = 273 pairs of photos in total). Three male rhesus macaques (M1: C, 19 years old, 13 kg; M2: F, 11 years old, 13 kg; M3: L, 20 years old, 11 kg) participated in the presidential election experiment, each for 5 days/sessions, with an intersession interval of at least 2 days in-between (n = 15 sessions in total). Stimulus sets for each session consisted of 13 pairs of presidential and vice-presidential candidate pictures.

### Experimental Setup

Each subject monkey sat in a primate chair (Crist Instruments), in a dark room (luminance ∼3 cd/m2) facing an LCD monitor (BenQ XL2730, 27’’, 2560*1440, 120 Hz). A computer (Dell Precision Tower 5810, custom built) running MATLAB (Mathworks) and Psychtoolbox (52, 53) was used to control all aspects of the experiment, including displaying visual stimuli on the monitor, communicating with the eye tracking system (Eyelink, see below), and opening and closing solenoid valves (Christ Instrument) to dispense juice rewards.

During the experiment, the monitor displayed a uniform gray background (luminance ∼15 cd/ m2). At the beginning of each trial, a central fixation spot (0.5°, luminance ∼35 cd/m2) was illuminated, and the monkey brought his gaze within a 3.0° diameter fixation window to initiate image display. Subsequently a pair of luminance-balanced, black-and-white images of competing political candidates were rendered on each side of the screen for 2.5 seconds. A blank screen then replaced both images, and a fixed amount of juice (0.5 ml) was delivered to the subject monkey. The inter-trial interval was 2-3 seconds (jittered). After initial fixation, the subject monkey was free to look anywhere during stimulus presentation as well as during the inter-trial interval. Eye position was recorded with an infrared eye tracking system, Eyelink 1000 Plus (SR Research, primate mount), sampled at 1,000 Hz, exported as EDF files, and then pre-processed with a custom MATLAB script (Edf2Mat, https://github.com/uzh/edf-converter).

### Stimulus presentation

Candidate photos for real U.S. gubernatorial, senatorial, and presidential general elections were presented in pairs. In each session, one set of races (gubernatorial, senatorial, or presidential) was presented in its entirety with each candidate pair displayed once and once only. The order (i.e. which race) and side (i.e. left or right side of the screen) of presentation were randomized.

#### Gubernatorial races

The procedure followed 18. From the *Almanac of American Politics*, a list was compiled of all gubernatorial races from 1995 to 2006, excluding races with highly familiar politicians (e.g., Arnold Schwarzenegger). There were 124 races (248 candidates) in total, with 36 female candidates, and 74 incumbents in running. Pictures of the winner and the runner-up were collected from various Internet sources (e.g., CNN, Wikipedia, and local media sources). The image of each politician was cropped, placed on a standard background, and converted to grayscale.

#### Senatorial races

The procedure followed 16, 18. From the *Almanac of American Politics*, a list was compiled of all senatorial races from 2000 to 2008, excluding races with highly familiar politicians (e.g., Hillary Clinton). There were 149 races (298 candidates) in total, with 44 female candidates, and 120 incumbents in running. Pictures of the winner and the runner-up were collected from various Internet sources (e.g., CNN, Wikipedia, and local media sources). The image of each politician was cropped, placed on a standard background, and converted to grayscale.

#### Presidential races

A list was compiled of all presidential races from 2000 to 2024, as well as the vice-presidential race of 2024. For 2000-2012, the most time-appropriate official portrait was chosen for each candidate. For example, for Obama vs Romney 2012, the 2012 presidential portrait of Obama and 2006 gubernatorial portrait of Romney were chosen. For more recent elections (2016-2024), since candidate age had become a major issue, in addition to the most time-appropriate official portraits, we also selected one specific election year photo for each candidate, from national conventions or major campaigning events. The image of each politician was cropped, placed on a standard background, and converted to grayscale.

### Data Analysis

Saccades and fixations were determined using Eyelink 1000 Plus online parser. All subsequent data analysis was done in custom MATLAB scripts. All statistical tests were two-tailed. For hypothesis testing between two samples, a non-parametric Wilcoxon signed rank test (for paired samples) or Wilcoxon rank sum test (for unpaired samples) was used. For comparison among more than two samples, an ANOVA was used controlling for multiple comparisons (Tukey’s HSD test) when appropriate. Correlation coefficients were estimated with Pearson’s r, or Spearman’s ρ when normality could not be assumed. Means were reported with standard errors of the mean (S.E.M.s); correlation coefficients were reported with 95% confidence intervals.

#### Competence/masculinity ratings

The procedure followed 16, 18.

#### Age

Age was calculated by subtracting the birth year of each candidate from the election year.

#### Facial feature measurements

The procedure followed 39. All measurements were performed in MATLAB Image Viewer. Jaw width, cheekbone width, face height and lower face height were measured in pixels, and normalized against inter-pupillary distance in pixels. Miscellaneous measurements included baldness (yes or no), facial hair (yes or no), glasses (yes or no), and smile (0: no smile; 1: closed-mouth smile; 2: open-mouth smile). At least two independent coders measured each face, and their ratings were averaged. The average concordance across raters was 0.79.

#### Red/blue /swing states

For each election, we used the presidential election closest in time to categorize each state as a Republican (i.e. red), Democratic (i.e. blue), or swing state. A swing state was defined as a state in which the difference in vote share between the Republican and Democratic presidential candidates was less than 10%. For example, for the 2008 senatorial election (Graham vs Conley), South Carolina was considered a swing state, as in the 2008 presidential election McCain won the state by a margin of 9%.

## Acknowledgments

We thank Dr. Carla Escabi, Dr. Leah Makaron, Dr. Kristin Gardiner, and the University Laboratory Animal Resources at University of Pennsylvania for providing excellent animal care. We thank Dr. Christopher Olivola and Dr. Gideon Nave for sharing the sets of gubernatorial and senatorial candidate photos and corresponding subject ratings. We thank Cajal, Frans, Oskar, and most of all Leakey, for participating in these experiments.

## Figure Legends

**Figure S1. a:** Example gaze patterns for gubernatorial elections in South Dakota (2006, top) and Alaska (2002, bottom). Cross: central fixation spot. Filled circles: fixations on candidate pictures; circle size: fixation duration. Magenta: fixations on election winner; green: fixations on loser. **b:** Gaze bias for all 3 male monkeys as a function of election outcome (top), and candidate gender (bottom). Error bars: mean ± SEM. **c:** Example gaze patterns in races female candidate prevailed over male opponent for gubernatorial elections in Alaska (2002, top) and Kansas (2006, bottom). Cross: central fixation spot. Filled circles: fixations on candidate pictures; circle size: fixation duration. Magenta: fixations on election winner; green: fixations on loser. **d:** Gaze bias as a function of election outcome for female vs male races. Error bars: mean ± SEM. **e:** Gaze bias election predictions across analytical approaches. **f:** Correlation between gaze bias and vote share restricted to races with incumbents. Error bars: 95% confidence interval.

**Figure S2. a:** Jawline bias as a function of election outcome. **b:** Jawline election outcome bias (loser vs. winner) as a function of election type (left) or winner identity (right). G-S: Gubernatorial and Senatorial races; FM-MM: Female-Male and Male-Male races; CC-CI: Challenger-Challenger and Challenger-Incumbent races; BS-RS: Blue State and Red State races; F-M win: Female-won and Male-won races; C-I win: Challenger-won and Incumbent-won races; O-Y win: Older-candidate-won and Younger-candidate-won races; D-R win: Democrat-won and Republican-won races. **c:** All facial masculinity cues as a function of candidate gender. **d:** All facial masculinity cues as a function of election outcome. Error bars: mean ± SEM. **e:** Correlation of jaw prominence with vote share for both male and female candidates. **f:** Correlations between all facial masculinity cues and vote share for female candidates. **g:** Correlation of monkey gaze bias with vote share as a function of candidate gender. **h:** Correlation between age and vote share. Error bars: 95% confidence interval.

**Figure S3. a:** Monkey gaze bias (loser vs winner) as a function of winner identity. F-M win: Female-won and Male-won races; O-Y win: Older-candidate-won and Younger-candidate-won races; C-I win: Challenger-won and Incumbent-won races; D-R win: Democrat-won and Republican-won races. **b:** Jawline bias (loser vs. winner) for blue, swing, and red states. Error bars: mean ± SEM. **c:** Correlation of monkey gaze bias with jaw prominence for blue, swing, and red states. Error bars: 95% confidence interval. **d:** All 3 male monkeys showed gaze biases towards the Democratic over the Republican presidential candidate. **e:** Gaze bias for elections between Donald Trump and Clinton, Biden, and Harris. Error bars: mean ± SEM. **f:** Trump’s jaw prominence vs. vote share in 2020 in relation to the overall correlation between the two (linear regression line).

## References

1. Quattrone, G. A., & Tversky, A. (1988). Contrasting rational and psychological analyses of political choice. American Political Science Review, 82(3), 719–736.

2. Zaller, J. (1992). The nature and origins of mass opinion. Cambridge University.

3. Kuklinski, J. H., & Quirk, P. J. (2000). Reconsidering the rational public: Cognition, heuristics, and mass opinion. Elements of reason: Cognition, choice, and the bounds of rationality, 153–182.

4. Cunningham, F. (2002). Theories of democracy: A critical introduction. Routledge.

5. Alvis, J. D., Blitz, M., Burns, T., Burns, D. E., Carrington, A. M., Clinton, D., … & Zuckert, C. H. (2021). Democracy and the History of Political Thought. Rowman & Littlefield.

6. Kahneman, D., & Tversky, A. (1984). Choices, values, and frames. American psychologist, 39(4), 341.

7. Kahneman, D., Slovic, P. & Tversky, A. (1982). Judgment under uncertainty: Heuristics and biases. Cambridge University Press.

8. Kahneman, D. (2003). Maps of bounded rationality: Psychology for behavioral economics. American economic review, 93(5), 1449–1475.

9. Smith, V. L. (2003). Constructivist and ecological rationality in economics. American economic review, 93(3), 465–508.

10. Glimcher, P. W. (2004). Decisions, uncertainty, and the brain: The science of neuroeconomics. MIT press.

11. Camerer, C., Loewenstein, G., & Prelec, D. (2005). Neuroeconomics: How neuroscience can inform economics. Journal of economic Literature, 43(1), 9–64.

12. Todd, P. M., & Gigerenzer, G. (2012). Ecological rationality: Intelligence in the world. OUP USA.

13. Gigerenzer, G., & Gaissmaier, W. (2011). Heuristic decision making. Annual review of psychology, 62(1), 451–482.

14. Platt, M.L. (2020). The leader’s brain. Wharton School Press.

15. Martin, D.S. (1978). Person perception and real-life electoral behavior. Australian Journal of Psychology, 30, 255.

16. Todorov, A., Mandisodza, A. N., Goren, A., & Hall, C. C. (2005). Inferences of competence from faces predict election outcomes. Science, 308(5728), 1623–1626.

17. Willis, J., & Todorov, A. (2006). First impressions: Making up your mind after a 100-ms exposure to a face. Psychological science, 17(7), 592–598.

18. Ballew, C. C., & Todorov, A. (2007). Predicting political elections from rapid and unreflective face judgments. Proceedings of the National Academy of Sciences, 104(46), 17948–17953.

19. Lawson, C., Lenz, G. S., Baker, A., & Myers, M. (2010). Looking like a winner: Candidate appearance and electoral success in new democracies. World Politics, 62(4), 561–593.

20. Poutvaara, P., Jordahl, H., & Berggren, N. (2009). Faces of politicians: Babyfacedness predicts inferred competence but not electoral success. Journal of experimental social psychology, 45(5), 1132–1135.

21. Antonakis, J., & Dalgas, O. (2009). Predicting elections: Child’s play!. Science, 323(5918), 1183–1183.

22. Olivola, C. Y., & Todorov, A. (2010). Elected in 100 milliseconds: Appearance-based trait inferences and voting. Journal of nonverbal behavior, 34, 83–110.

23. Abelson, R. P., Kinder, D. R., Peters, M. D., & Fiske, S. T. (1982). Affective and semantic components in political person perception. Journal of personality and social psychology, 42(4), 619.

24. Zebrowitz, L. (2018). Reading faces: Window to the soul?. Routledge.

25. Waitt, C., & Little, A. C. (2006). Preferences for symmetry in conspecific facial shape among Macaca mulatta. International Journal of Primatology, 27, 133–145.

26. Costa, M., Gomez, A., Barat, E., Lio, G., Duhamel, J. R., & Sirigu, A. (2018). Implicit preference for human trustworthy faces in macaque monkeys. Nature Communications, 9(1), 4529.

27. Deaner, R.O., Khera, A.V., Platt, M.L. (2005). Monkeys pay per view: adaptive valuation of social images by rhesus macaques. Current Biology. 15(6):543–548.

28. Watson, K.K., Ghodasra, J.H., Furlong, M.A., Platt, M.L. 2012. Visual preferences for sex and status in female rhesus macaques. Animal cognition. 15(3):401–407.

29. Shepherd, S. V., Deaner, R. O., & Platt, M. L. (2006). Social status gates social attention in monkeys. Current biology, 16(4), R119–R120.

30. Ebitz, R. B., Watson, K. K., & Platt, M. L. (2013). Oxytocin blunts social vigilance in the rhesus macaque. Proceedings of the National Academy of Sciences, 110(28), 11630–11635.

31. Jiang, Y., Sheng, F., Belkaya, N., & Platt, M. L. (2022). Oxytocin and testosterone administration amplify viewing preferences for sexual images in male rhesus macaques. Philosophical Transactions of the Royal Society B, 377(1858), 20210133.

32. Klein, J. T., Shepherd, S. V., & Platt, M. L. (2009). Social attention and the brain. Current Biology, 19(20), R958–R962.

33. Cover, A. D. (1977). One good term deserves another: The advantage of incumbency in congressional elections. American Journal of Political Science, 523–541.

34. Gelman, A., & King, G. (1990). Estimating incumbency advantage without bias. American journal of political science, 1142–1164.

35. Klein, J. T., Deaner, R. O., & Platt, M. L. (2008). Neural correlates of social target value in macaque parietal cortex. Current Biology, 18(6), 419–424.

36. Klein, J. T., & Platt, M. L. (2013). Social information signaling by neurons in primate striatum. Current Biology, 23(8), 691–696.

37. Smuts, B. B., Cheney, D. L., Seyfarth, R. M., & Wrangham, R. W. (Eds.). (2008). Primate societies. University of Chicago Press.

38. Grammer, K., & Thornhill, R. (1994). Human (Homo sapiens) facial attractiveness and sexual selection: the role of symmetry and averageness. Journal of comparative psychology, 108(3), 233.

39. Penton-Voak, I. S., Jones, B. C., Little, A. C., Baker, S., Tiddeman, B., Burt, D. M., & Perrett, D. I. (2001). Symmetry, sexual dimorphism in facial proportions and male facial attractiveness. Proceedings of the Royal Society of London. Series B: Biological Sciences, 268(1476), 1617–1623.

40. Rhodes, G. (2006). The evolutionary psychology of facial beauty. Annual Review in Psychology, 57(1), 199–226.

41. Bartels, L. M. (2000). Partisanship and voting behavior, 1952–1996. American Journal of Political Science, 44, 35–50.

42. Waitt, C., Little, A. C., Wolfensohn, S., Honess, P., Brown, A. P., Buchanan-Smith, H. M., & Perrett, D. I. (2003). Evidence from rhesus macaques suggests that male coloration plays a role in female primate mate choice. Proceedings of the Royal Society of London. Series B: Biological Sciences, 270(uppl_2), S144–S146.

43. Dubuc, C., Allen, W. L., Maestripieri, D., & Higham, J. P. (2014). Is male rhesus macaque red color ornamentation attractive to females?. Behavioral ecology and sociobiology, 68, 1215–1224.

44. Keating, C. F. (1985). Gender and the physiognomy of dominance and attractiveness. Social psychology quarterly, 61–70.

45. Scheib, J. E., Gangestad, S. W., & Thornhill, R. (1999). Facial attractiveness, symmetry and cues of good genes. Proceedings of the Royal Society of London. Series B: Biological Sciences, 266(1431), 1913–1917.

46. Mazur, A., Mazur, J., & Keating, C. (1984). Military rank attainment of a West Point class: Effects of cadets’ physical features. American Journal of Sociology, 90(1), 125–150.

47. Pearson, J. M., Watson, K. K., & Platt, M. L. (2014). Decision making: the neuroethological turn. Neuron, 82(5), 950–965.

48. Platt, M. L., Seyfarth, R. M., & Cheney, D. L. (2016). Adaptations for social cognition in the primate brain. Philosophical Transactions of the Royal Society B: Biological Sciences, 371(1687), 20150096.

49. Kahn, K. F., & Kenney, P. J. (2002). The slant of the news. American Political Science Review, 96, 381– 394.

50. Barrett, A. W., & Barrington, L. W. (2005b). Bias in newspaper photograph selection. Political Research Quarterly, 58, 609–618.

51. Little, A. C., Burriss, R. P., Jones, B. C., & Roberts, S. C. (2007). Facial appearance affects voting decisions. Evolution and Human Behavior, 28(1), 18–27.

52. Brainard, D. H., & Vision, S. (1997). The psychophysics toolbox. Spatial vision, 10(4), 433–436.

53. Pelli, D. G. (1997). The VideoToolbox software for visual psychophysics: transforming numbers into movies. Spatial vision, 10(4), 437–442.

