## Supplementary figures and images for "Monkeys Predict US Elections"

### Supplementary Figures

#### Supplementary Figure 1

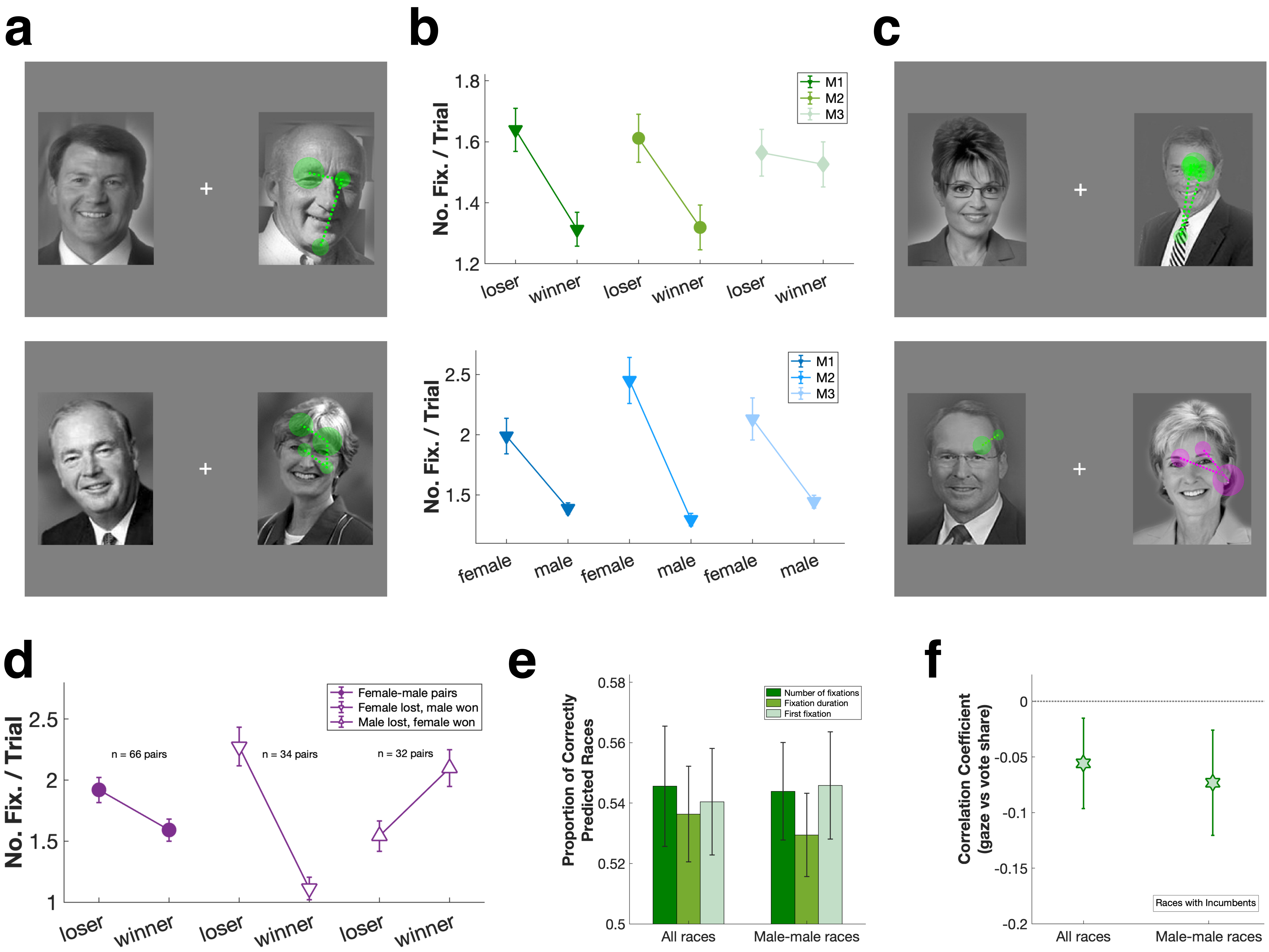

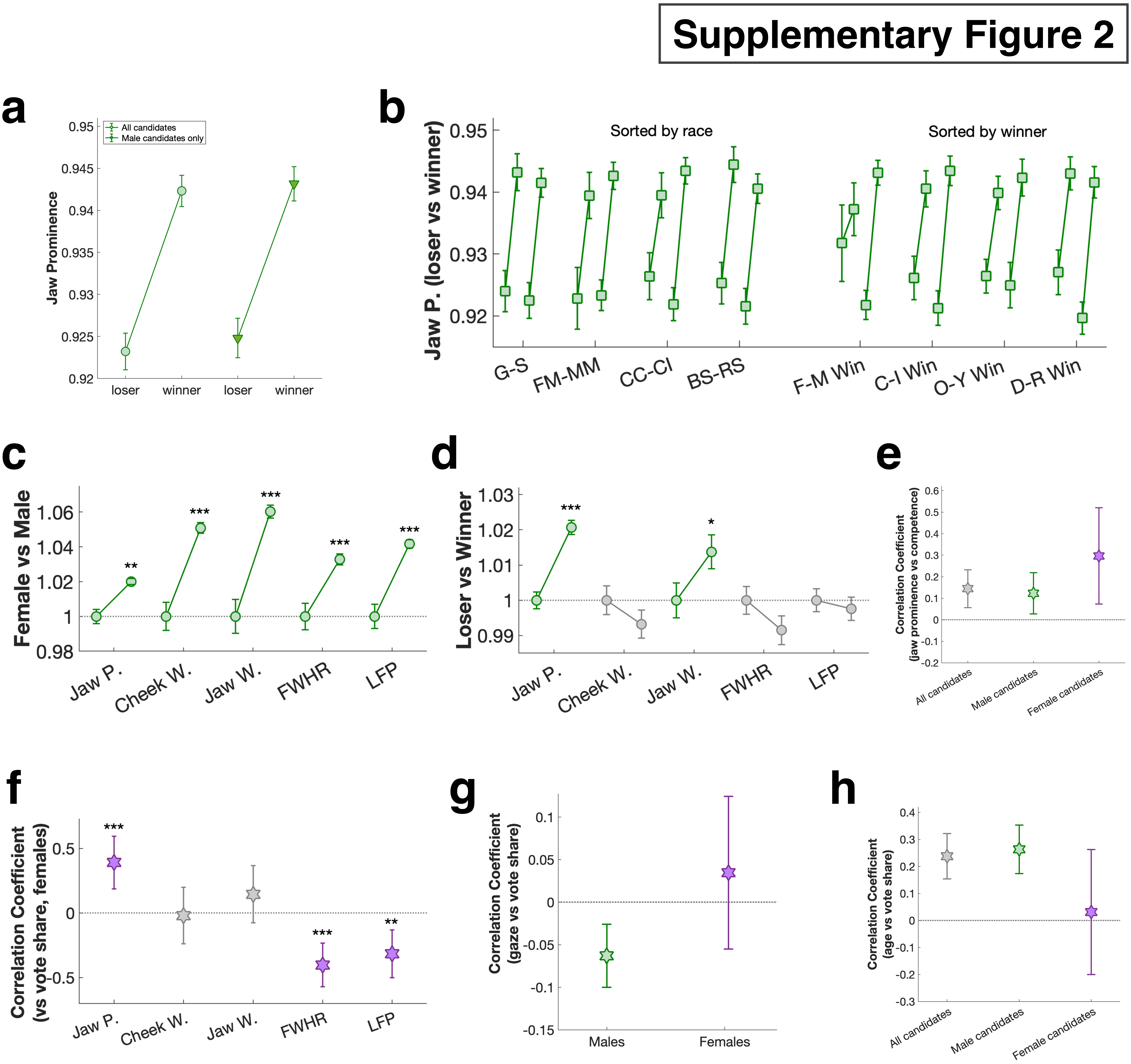

##### Supplementary Figure 3

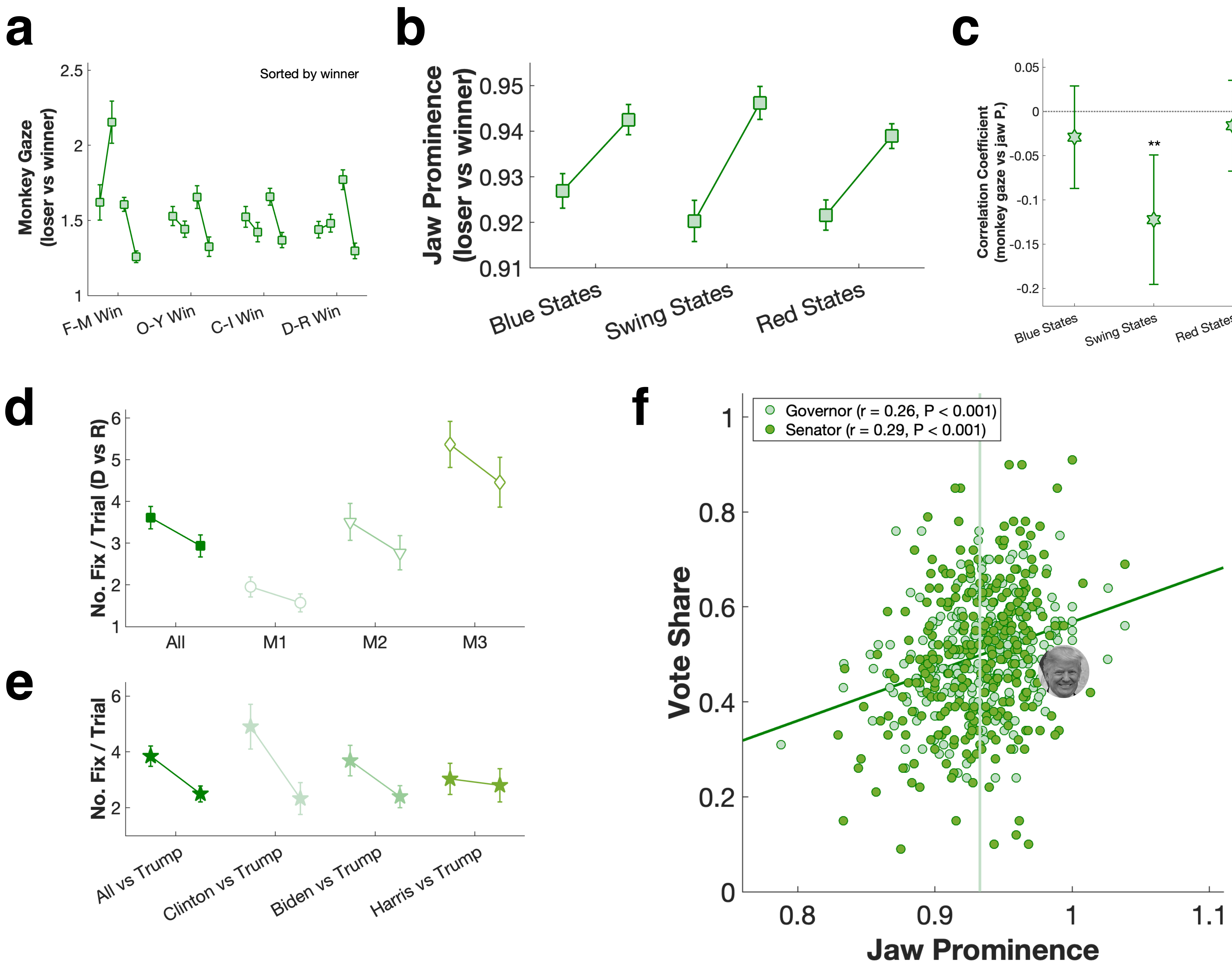
